# Integrative Structural Brain Network Analysis In Diffusion Tensor Imaging

**DOI:** 10.1101/129015

**Authors:** Moo K. Chung, Jamie L. Hanson, Nagesh Adluru, Andrew L. Alexander, Richard J. Davidson, Seth D. Pollak

## Abstract

In diffusion tensor imaging, structural connectivity between brain regions is often measured by the number of white matter fiber tracts connecting them. Other features such as the length of tracts or fractional anisotropy (FA) are also used in measuring the strength of connectivity. In this study, we investigated the effects of incorporating the number of tracts, the tract length and FA-values into the connectivity model. Using various node-degree based graph theory features, the three connectivity models are compared. The methods are applied in characterizing structural networks between normal controls and maltreated children, who experienced maltreatment while living in post-institutional settings before being adopted by families in the US.

## 1 Introduction

Diffusion tensor imaging (DTI) is a non-invasive imaging modality often used to characterize the microstructure of biological tissues using magnitude, anisotropy and anisotropic orientation associated with diffusion (Basser et al., 1994). It is assumed that the direction of greatest diffusivity is most likely aligned to the local orientation of the white matter fibers. Traditionally scalar measures such as fractional anisotropy (FA) and mean diffusivity (MD) obtained from DTI have been used for quantifying clinical populations at the voxel-level (Basser and Pierpaoli, 1996; Barnea-Goraly et al., 2004; Roberts et al., 2005; Jones et al., 2006; Smith et al., 2006; Daianu et al., 2013). Various tractography methods have been developed to visualize and map out major white matter pathways in individuals and brain atlases (Basser et al., 2000; Catani et al., 2002; Conturo et al., 1999; Lazar et al., 2003; Mori et al., 1999, 2002; Thottakara et al., 2006; Yushkevich et al., 2007). The tractography can yield additional connectivity metrics that describe the localized variations in connectivity strength as a form of network graphs. The tractography based whole brain network analysis has shown considerable promise quantifying neural pathways in various populations (Daianu et al., 2013).

The strength of connection from one gray matter region to another is often measured by counting the number of fiber tracts connecting the two regions in predefined parcellations (Daianu et al., 2013; Gong et al., 2009). A problem with the simple tract counting approach might be that the neglect of both the distance between the regions and FA-values. There have been few modifications and variations to the simple tract counting method in the literature. Skudlarski et al. (2008) used a weighting scheme that penalizes indirect longer connections. The connectivity between two parcellations is given by tract counts normalized by the volume of region of interest (Van Den Heuvel and Sporns, 2011). Kim et al. (2015) and Van Den Heuvel and Sporns (2011) used the mean FA-values along the tracts as the measure of connectivity. However, it is unclear if there is an optimal connectivity measure, or if different approaches meaningfully impact results.

In this study, connectivity methods based on tract count, length and its FA-values are compared. The tract length and its FA-value based methods are using an electrical circuit model as an analogy, where the strength of connection corresponds to the resistance of the circuit. Although electrical circuit models were never used for modeling brain networks at the macroscopic level except Chung (2012) and Chung et al. (2012), they were often used in a wide variety of mostly biological networks that are not related to any electrical circuit. Starting with Doyle and Snell (1984), numerous studies formulated graphs and networks as electrical circuits (Chandra et al., 1996; Tetali, 1991). Segev et al. (1985) used an electrical circuit model to model the electrical behavior of neurons. Electrical resistance models are also used in genetic networks (Leiserson et al.; Suthram et al., 2008). In particular, Yeger-Lotem et al. (2009) and Basha et al. (2013) formulated a minimum-cost network flow in genetic networks, which is related to electrical resistance.

For comparisons between the three methods, graph theoretical features based on node degree is investigated. The node degree, which counts the number of connections at a node, is probably the most fundamental graph theoretic feature used in network analysis. The node degree is used directly and indirectly defining many other graph theoretic features (Bullmore and Sporns, 2009; Fornito et al., 2016). The node degree and its probability distribution will be investigated in detail. To complement the comparisons, the strength of connectivity will be also contrasted. The comparisons will be done by contrasting two different samples: normal controls and maltreated children, who experienced severe early life stress and maltreatment.

Early and severe childhood stress, such as experiences of abuse and neglect, have been associated with a range of cognitive deficits (Pollak, 2008; Sanchez and Pollak, 2009; Loman et al., 2010) and structural abnormalities (Jackowski et al., 2009; Hanson et al., 2012, 2013; Gorka et al., 2014) years following the stressors. However, little is known about the underlying biological mechanisms leading to cognitive problems in these children (Pollak et al., 2010). Four studies have reported alterations in prefrontal white matter in children exposed to early stress (De Bellis et al., 2002; Hanson et al., 2012, 2013, 2015). Other studies have suggested that early stress may cause larger hippocampal white matter volume (Tupler and De Bellis, 2006) or smaller cerebellar volume (Bauer et al., 2009). The extant literature has been based upon region of interest (ROI) based volumetry or univariate vowel-wise morphometric techniques to characterize anatomical differences between groups. Network approaches may allow for a richer and fuller characterization of specific neural circuits, facilitating greater understanding in brain and behavioral alterations after child maltreatment. In this study, we describe three network analysis frameworks and demonstrate the potential utility by applying it to DTI comparisons of children exposed to early life stress and maltreatment.

## 2 Methods

### 2.1 Subjects

The study consisted of 23 children who experienced documented maltreatment early in their lives, and 31 age-matched normal control (NC) subjects. All the children were recruited and screened at the University of Wisconsin-Madison. The maltreated children were raised in institutional settings, where the quality of care has been documented as falling well below the standard necessary for healthy human development. For the controls, we selected children without a history of maltreatment from families with similar current socioeconomic statuses. The exclusion criteria included, among many others, abnormal IQ (< 78), congenital abnormalities (e.g., Down syndrome or cerebral palsy) and fetal alcohol syndrome (FAS). The average age for maltreated children was 11.26 ± 1.71 years while that of controls was 11.58 ± 1.61 years. This particular age range was selected since this development period is characterized by major regressive and progressive brain changes (Lenroot and Giedd, 2006; Hanson et al., 2013). There were 10 boys and 13 girls in the maltreated group and 18 boys and 13 girls in the control group. Groups did not differ on age, pubertal stage, sex, or socio-economic status (Hanson et al., 2013). The average amount of time spent in institutional care by children was 2.5 years ± 1.4 years, _with a range from 3 months to 5.4 years. Children were on average 3.2 years_ ± 1.9 months when they were adopted, with a range of 3 months to 7.7 years. Table 1 summarizes the participant characteristics. Additional details about the recruitment strategy and participants characteristics can be found in Hanson et al. (2013).

**Table 1:** Study participant characteristics

|  | Maltreated | Normal controls | Combined |
| --- | --- | --- | --- |
| Sample size | 23 | 31 | 54 |
| Sex (males) | 10 | 18 |  |
| Age (years) | $11.26 \pm 1.71$ | $11.58 \pm 1.61$ | |
| Duration (years) | $2.5 \pm 1.4$ (0.25 to 5.4) | | |
| Time of adoption (years) | $3.2 \pm 1.9$ (0.25 to 7.7) | | |

### 2.2 Preprocessing

DTI were collected on a 3T General Electric SIGNA scanner (Waukesha, WI) using a cardiac-gated, diffusion-weighted, spin-echo, single-shot, EPI pulse sequence. Diffusion tensor encoding was achieved using 12 optimum non-collinear encoding directions with a diffusion weighting of 1114 s/mm^2^ and a non-DW T2-weighted reference image. Other imaging parameters were TE = 78.2 ms, 3 averages (NEX: magnitude averaging), and an image acquisition matrix of 120 × 120 over a field of view of 240 × 240 mm^2^. The details on other image acquisition parameters are given in Hanson et al. (2013). The acquired voxel size of 2 × 2 × 3 mm was interpolated to 0.9375 mm isotropic dimensions (256 × 256 in plane image matrix) on the scanner during the image reconstruction using the zero-filled interpolation. This is a loss-less interpolation and there is no added blurring. The scanner setting was used in our previous studies (Chung et al., 2015; Hanson et al., 2013). To minimize field inhomogeneity and image artifacts, high order shimming and fieldmap images were collected using a pair of non-EPI gradient echo images at two echo times: TE1 = 8 ms and TE2 = 11 ms.

DTI processing follows the pipeline established in the previous DTI studies (Hanson et al., 2013; Kim et al., 2015). DTI were corrected for eddy current related distortion and head motion via FSL software (http://www.fmrib.ox.ac.uk/fsl) and distortions from field inhomogeneities were corrected using custom software based on the method given in Jezzard and Clare (1999) before performing a non-linear tensor estimation using CAMINO (Cook et al., 2006). Spatial normalization of DTI data was done using a diffeomorphic registration strategy (Joshi et al., 2004; Zhang et al., 2007) DTI-ToolKit (DTI-TK; http://www.nitrc.org/projects/dtitk). A population specific tensor template was constructed. Fractional anisotropy (FA) were calculated for diffusion tensor volumes diffeomorphically registered to the study specific template. Tractography was done in the normalized space using the TEND algorithm and warped into the study template (Lazaret al., 2003). We used Anatomical Automatic Labeling (AAL) with 116 parcellations (Tzourio-Mazoyer et al., 2002) (Figure 1). The AAL atlas was warped to the study template via the diffeomorphic image registration. The two end points of fiber tracts are identified with respect to 116 parcellations and the tract lengths are computed. Any tracts that do not pass through two given parcellations are removed. Tracts passing through only one parcellation are also removed.

**Figure 1:**
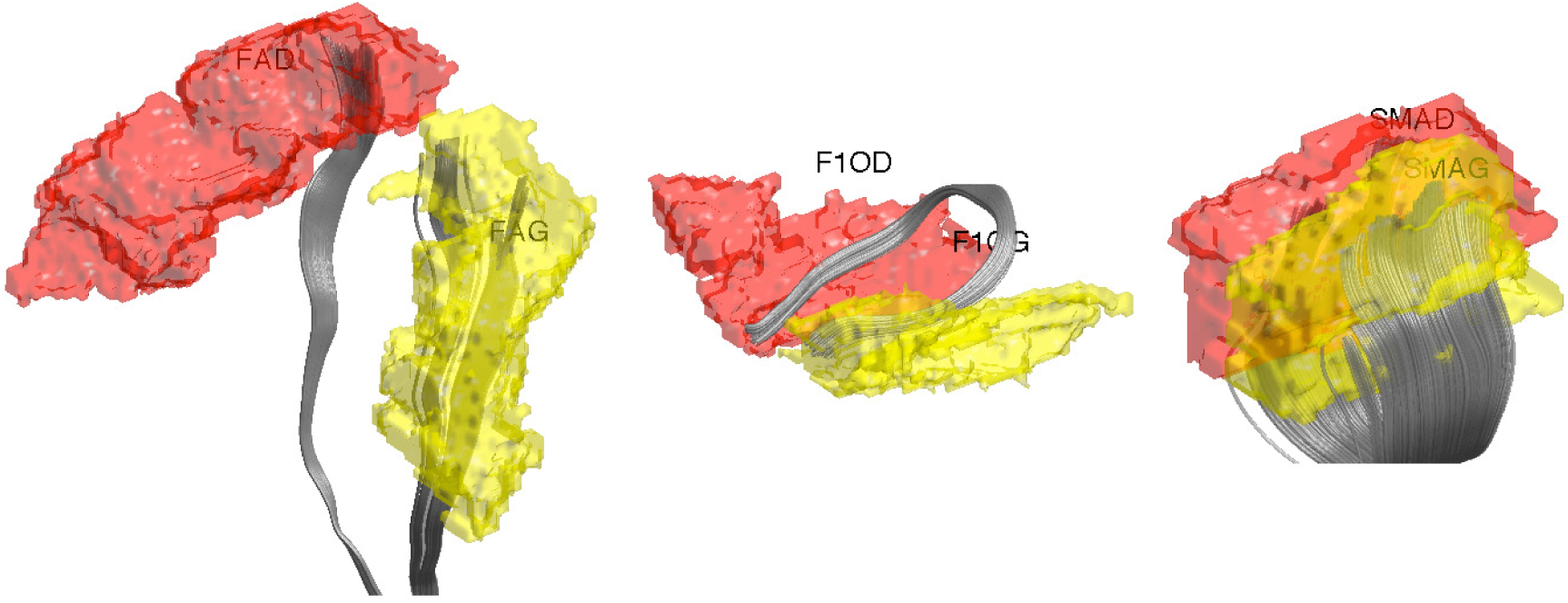
White matter fiber tracts connecting two regions in Anatomical Automatic Labeling (AAL) with 116 parcellations. The labels for parcellations are FAG (left pre-central) FAD (right precentral), F1OF (left frontal mid orbital), F1OG (right frontal mid orbital), SMAG (left superior motor area) and SMAD (right superior motor area).

### 2.3 Structural Connectivity Matrices

In order to have more integrative method for dealing with tract counts and lengths, we start with the arithmetic and harmonic means. Given measurements *R*_1_,…, *R_k_*, their arithmetic mean *A*(*R*_1_,…,*R_k_*) is given by the usual sample mean, i.e.,

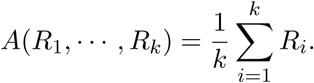

The tract count and its mean are based on the arithmetic addition. The *harmonic mean H*(*R*_1_;…;*R*_*k*_) of *R*_1_;…;*R_k_* is given by

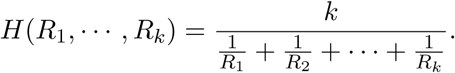

The harmonic mean is given by the reciprocal of the arithmetic mean of reciprocal of measurements:

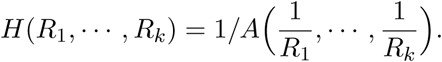

The harmonic mean has been mainly used in measuring the rates of a physical systems such as the speed a car (Zhang et al., 1999), resistance of a electrical circuits (?). Beyond physical systems, it has been used in k-means clustering (Zhang et al., 1999), where the harmonic k-mean is used instead of the usual arithmetic mean. The harmonic mean has been also used in the integrated likelihood for the Bayesian model selection problem (Raftery et al., 2006). Whenever we deal with rates and ratio based measures such as resistance, the harmonic mean provides more robust and accurate average compared to the arithmetic mean and often used in various branch of sciences. The use of harmonic means can naturally incorporate the length of tracts in connectivity.

Motivated by Doyle and Snell (1984), the DTI brain network can be analogously modeled as an electrical system consisting of series and parallel circuits (Figure 2). Each fiber tract may be viewed as a single wire with resistance *R* proportional to the length of the wire. If two regions are connected through an intermediate region, it forms a series circuit. In the series circuit, the total resistance *R* is additive so we have

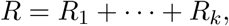

where *R*_*k*_ is the resistance of the *k*-th tract. If multiple fiber tracts connect two regions, it forms a parallel circuit, where the total resistance is

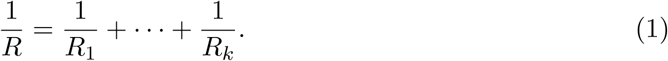

**Figure 2:**
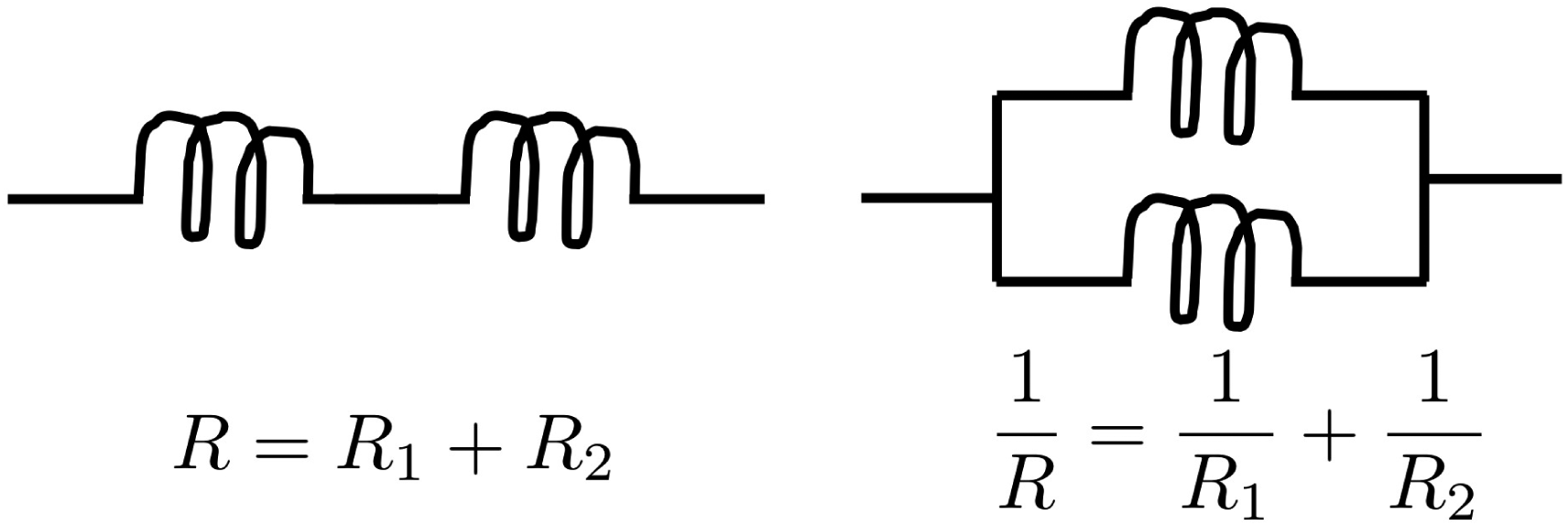
Left: series circuit. Right: parallel circuit.

The total resistance of a series circuit is related to the arithmetic mean of tract lengths. On the other hand, the total resistance (1) of a parallel circuit is related to the harmonic mean. Any complex parallel circuits in an electrical system can be simplified using a single wire with the equivalent resistance. Hence, we can simplify whole brain fiber tracts into smaller number of equivalent tracts in the model. The reciprocal of the resistance is then taken as the measure of connectivity. Smaller resistance corresponds to stronger connectivity. Figure 3 shows examples of parallel circuits. If all the tracts are 10 cm in length, the total resistance becomes 10, 5 and 2 as the number of tracts increases to 1, 2 and 5. The corresponding connectivities between A and B are 0.1, 0.2 and 0.5. Thus, if the tract lengths are all the same, the resistance based connectivity is proportional to tract counts. Figure 4 shows various toy networks and the corresponding resistance matrices. The corresponding resistance matrices given below.

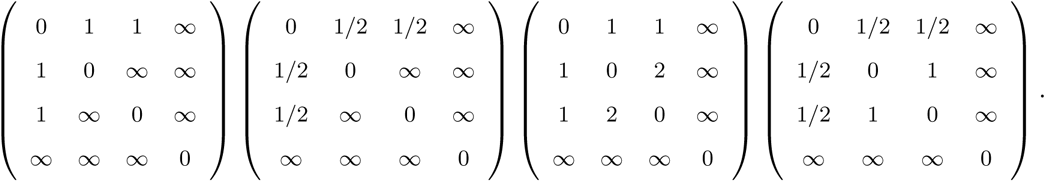

The resistance between indirectly connected nodes is *∞*. The most redundant network has the smallest resistance. The reciprocal of the resistance is taken as the strength of connectivity.

**Figure 3:**
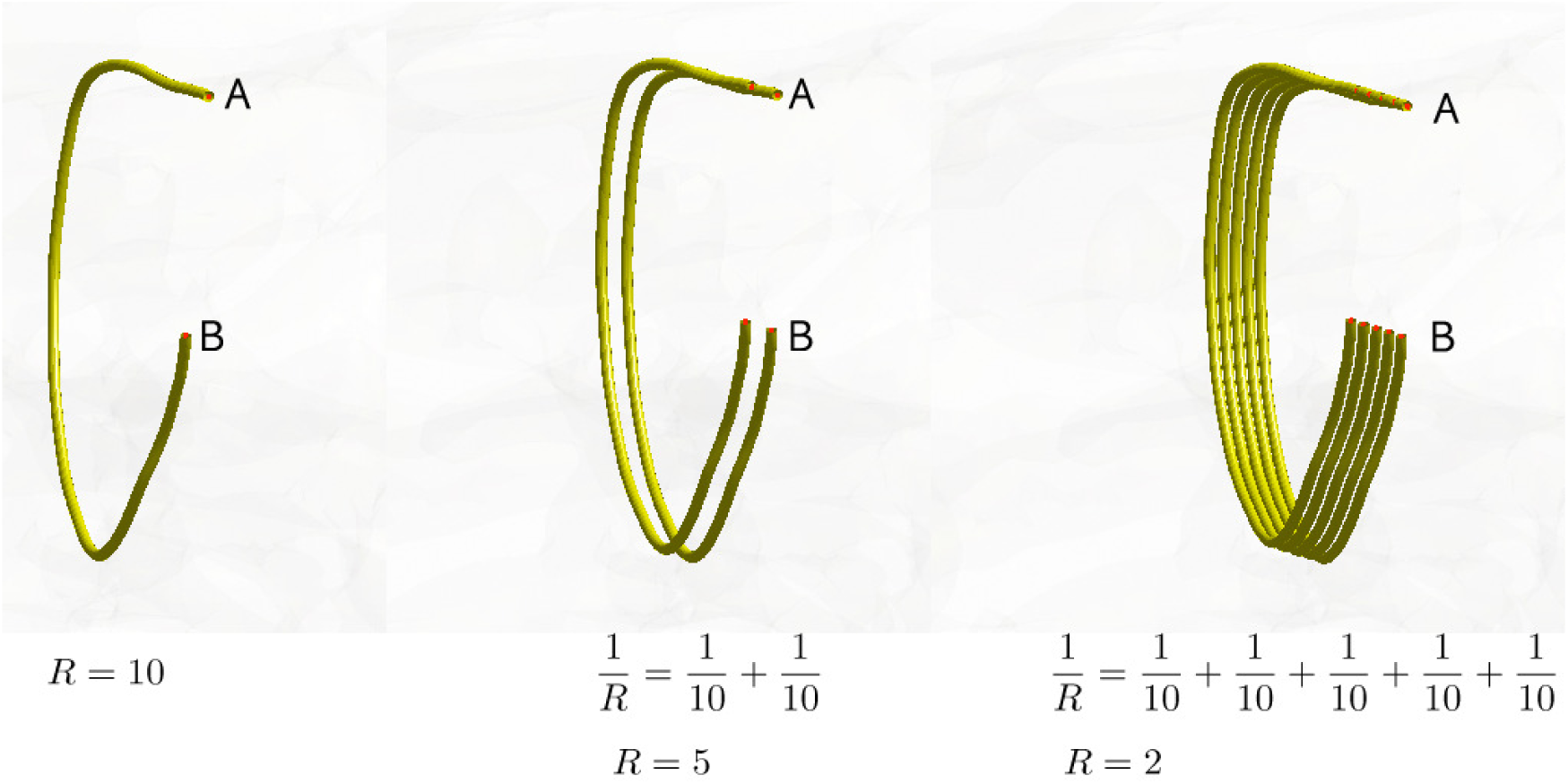
Multiple fiber tracts connecting the regions A and B are modeled as a parallel circuit. The resistance in a wire is proportional to the length of the wire. As more tracts connect the regions in parallel, the resistance decreases and the strength of connection increases. In this example, we let the resistance of each tract equal to the length of the tract. If all the tracts are 10 cm in length, the total resistance becomes 10, 5 and 2 as the number of tracts increases to 1, 2 and 5. The connectivity between A and B is defined as the reciprocal of resistance. The corresponding connectivities are 0.1, 0.2 and 0.5.

**Figure 4:**
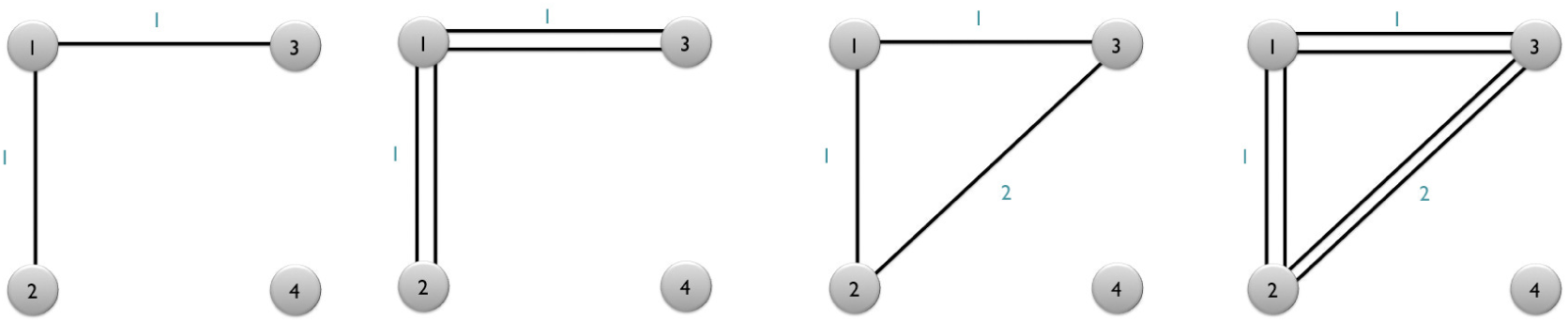
Toy networks with different connectivity resistance. The numbers between nodes are the length of tracts.

The electronic circuit model can be used in constructing a simplified but equivalent brain network. The two end points of tracts are identified. All the parallel tracts between any two regions are identified and replaced with a single tract with the equivalent resistance. This process completely removes all the parallel circuits. At the end, the simplified circuit forms a graph with the resistances as the edge weights.

Consider a tract 𝓜 consisting of *n* control points *p*_1_,…, *p_n_* obtained through tractography algorithms. Consider an inverse map ς^-1^ hat maps the control point *P_j_* onto the unit interval as

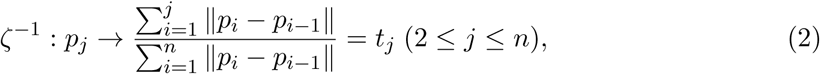

where ||. || is the Euclidean distance. This is the ratio of the arc-length from the point *p*_1_ to *p*_*j*_, to *p*_1_ to *P*_*n*_. We let this ratio be *t*_*j*_. We assume ς^-1^(*p*_l_) = 0. The ordering of the control points is also required in obtaining smooth one-to-one mapping. We parameterize the tract on a unit interval:

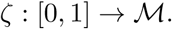

Then the total length *L*(𝓜) of the tract 𝓜 is given by

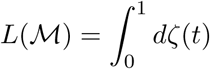

and discretely approximated as

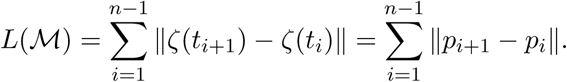

The connectivity matrices are subsequently determined by computing the resistance between parcellations (Figure 5-middle). We took the reciprocal ofthe resistance between the nodes as entries of the connectivity matrix. Smaller resistance corresponds to stronger connection. The method replaces a collection of parallel circuits with a single equivalent circuit. This process completely removes all the parallel circuits and simplifies complex parallel circuits to a simple circuit. The simplified circuit naturally forms a 3D graph with the resistances as the edge weights. Figure 6-middle shows the mean connectivities using the graph representation.

**Figure 5:**
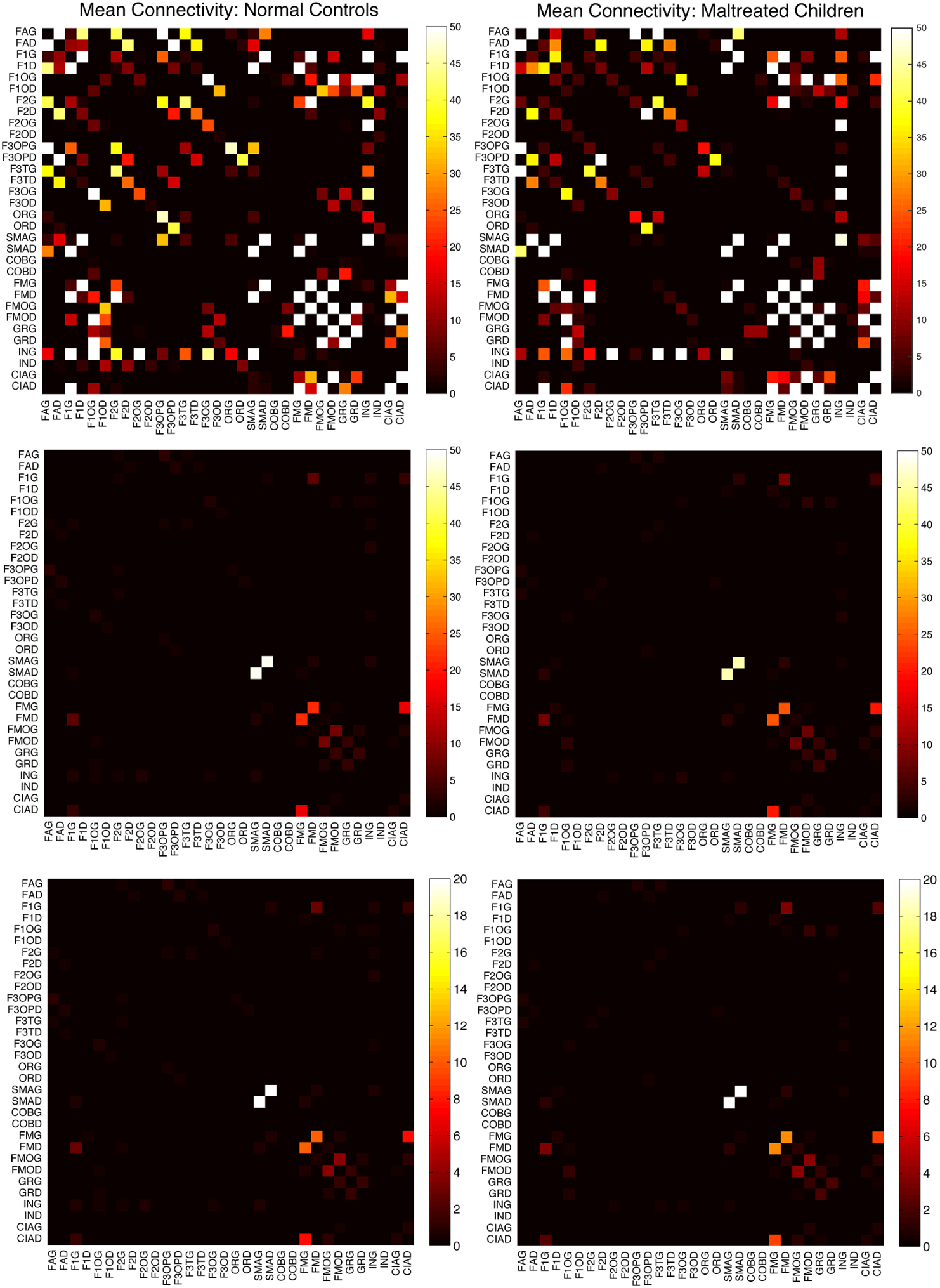
Top: mean connectivity matrices for first 32 nodes for normal controls (left) and maltreated children (right) using tract counts. Middle: the proposed electrical resistance based connectivity matrices. Although there are connectivity differences, strong connections are consistently shown in the two populations indicting the robust nature of the processing pipelines. The highest connectivity is shown between SMAG (left superior motor area) and SMAD (right superior motor area) in the both groups. Bottom: Resistance-based mean connectivity matrices that incorporate FA-values.

**Figure 6:**
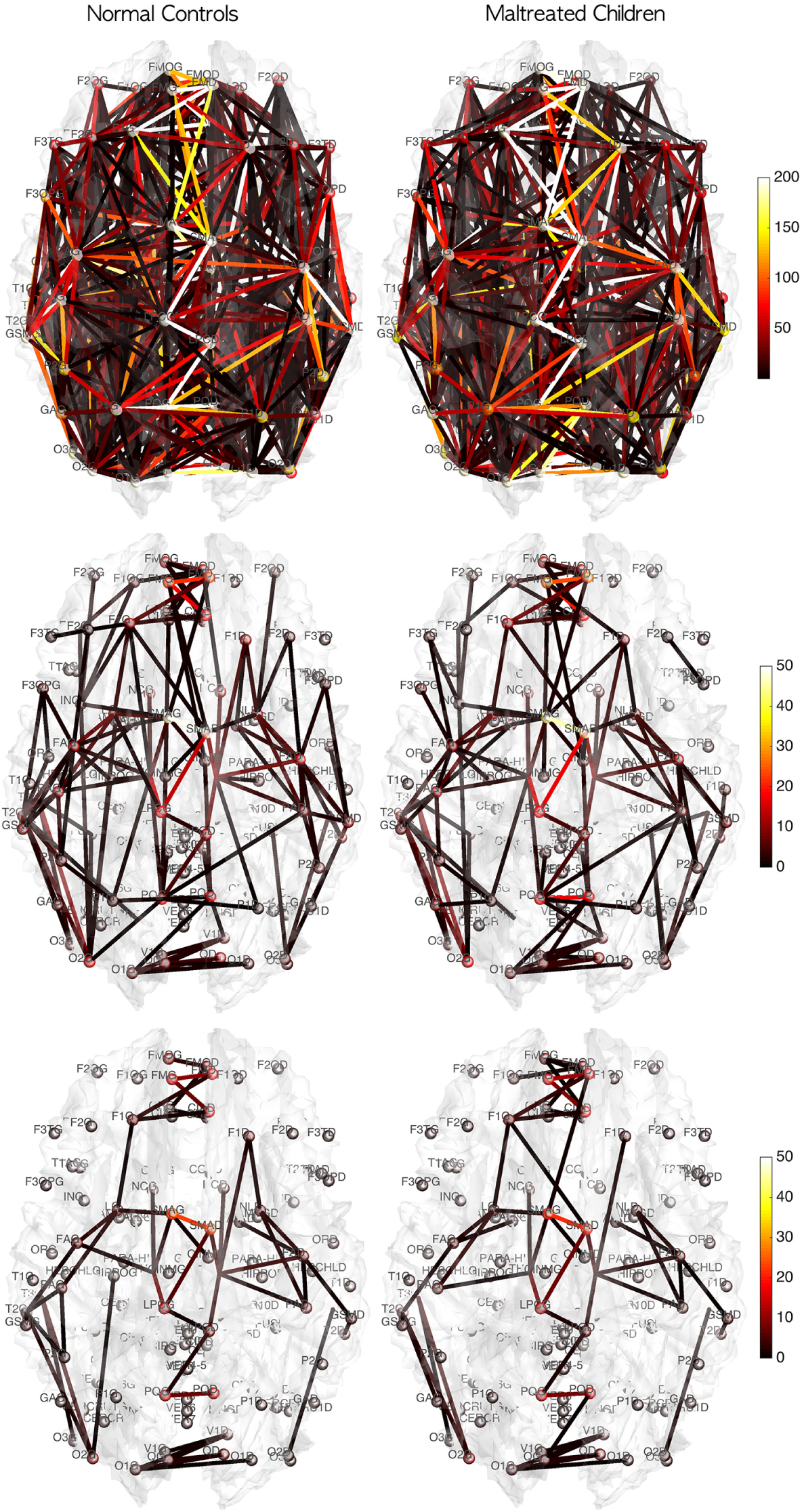
Mean connectivity of normal controls and maltreated children in the graph representation. The color of nodes and edges correspond to the connectivity strength in terms of tract counts (top), electrical resistance (middle), electrical resistance with FA-values (bottom). Although there are connectivity differences, strong connections are consistently shown in all three models.

Fractional anisotropy (FA) can be also used in constructing the connectivity matrix. FA may provide additional structural information that the tractography alone may not provide. Note that most parcellations are located in the gray matter regions, where the fiber tracts starts and ends so FA-values are expected to be very low at the nodes. The mass center of each parcellation is taken as a node of the network. The mean FA value at the node positions is 0.18 ± 0.10. In fact, at each node position, the two-sample *t*-statistic did not yield any group differences at 0.05 significance. However, along the tracts, it is expected FA-values are higher and it may influence the connectivity. Figure 8 shows an example of how FA-values change along 108 tracts between the left superior motor area (SMAG) to the right superior motor area (SMAD).

We incorporated FA-values along the fiber tracts as follows. By identifying voxels that tracts are passing through, we were able to linearly interpolate the FA-values along the tracts. If the FA value is larger at a certain part of a tract, it is more likely that the segment of the tract has been more stably estimated. Thus, the resistance of a tract segment can be modeled as inversely proportional to the FA value but proportional to the length of the segment *dζ* at point:

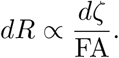

Note that we are using the reciprocal of the resistance as a tract connectivity metric *C*,
i.e.,

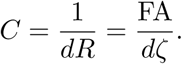

So heuristically, larger the value of FA, stronger the connectivity in the tract segment.

Subsequently, the total resistance *R* of the whole whole tract is defined as

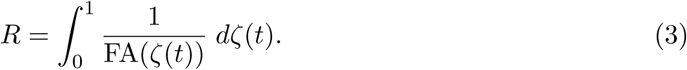

If the tractography processing is properly done, it is not possible to have zero FA-value along the obtained tracts so the integral (3) is well defined. The integral (3) is discretized as

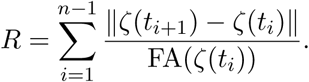

In our study, the step size ||*ζ*(*t*_*i*+1_) − *ζ*(*t_i_*)|| is fixed at 0.1 mm. Thus,

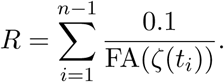

This is the weighted version of the resistance that incorporates FA-values. The connectivity matrices are then similarly determined by computing the reciprocal of the weighted resistance between parcellations (Figure 5-bottom). Smaller resistance corresponds to stronger connection. The pattern of connectivity matrices are almost identical to the connectivity without FA-values although the scale and local variations differ slightly.

### 2.4 How tract counts and length based connectivity are related

So far we presented three methods for constructing connectivity matrices in DTI. However, it is unclear how they are related and if they will give consistent statistical results at the end. Suppose there are *k* tracts between two parcellations. Suppose the tract lengths are all identical as *L*. Then the tract count based connectivity gives the connectivity strength *k*. The length based connectivity gives the connectivity strength *k/L*.

Under the ideal situation of the same tract length, the tract length based connectivity is proportional to tract count based connectivity. For the incorporation of FA-values to the connectivity, if we assume the identical FA-values along the tract, we have FA *· k/L* as connectivity. All three methods are proportional to each other. Thus, in the ideal situation of the same tract length and FA-values, the final statistical analyses for three methods would be identical since most statistical test procedures are scale invariant.

The difference arises in practice where the tract lengths are all different. This is a difficult problem we can’t address theoretically. To begin to deal with this issue, we checked the robustness of the tract length based method in relation to the tract count method. To determine the robustness of the tract length based metric, we calculated the relative error in comparison to traditional tract count based connectivity in the tract mislabeling problem.

Figure 7 shows a schematic of a possible tract mislabeling problem. Assume *k* number of tracts are passing through between parcellations 2 and 3 (top). In the traditional connectivity metric, tract count is used as the strength of connectivity. So the expected connectivity is *C*_*E*_ = *k*. However, *m* number of tracts might be mislabeled to pass through parcellations 1 and 3 (bottom). So the observed connectivity is given by *C*_*O*_ = *k*−*m*. The relative error in the tract count based connectivity is then (*C_E_* − *C_O_*)*/C_E_* = *m/k*. Let us determine the relative error for the tract length based connectivity metric.

**Figure 7:**
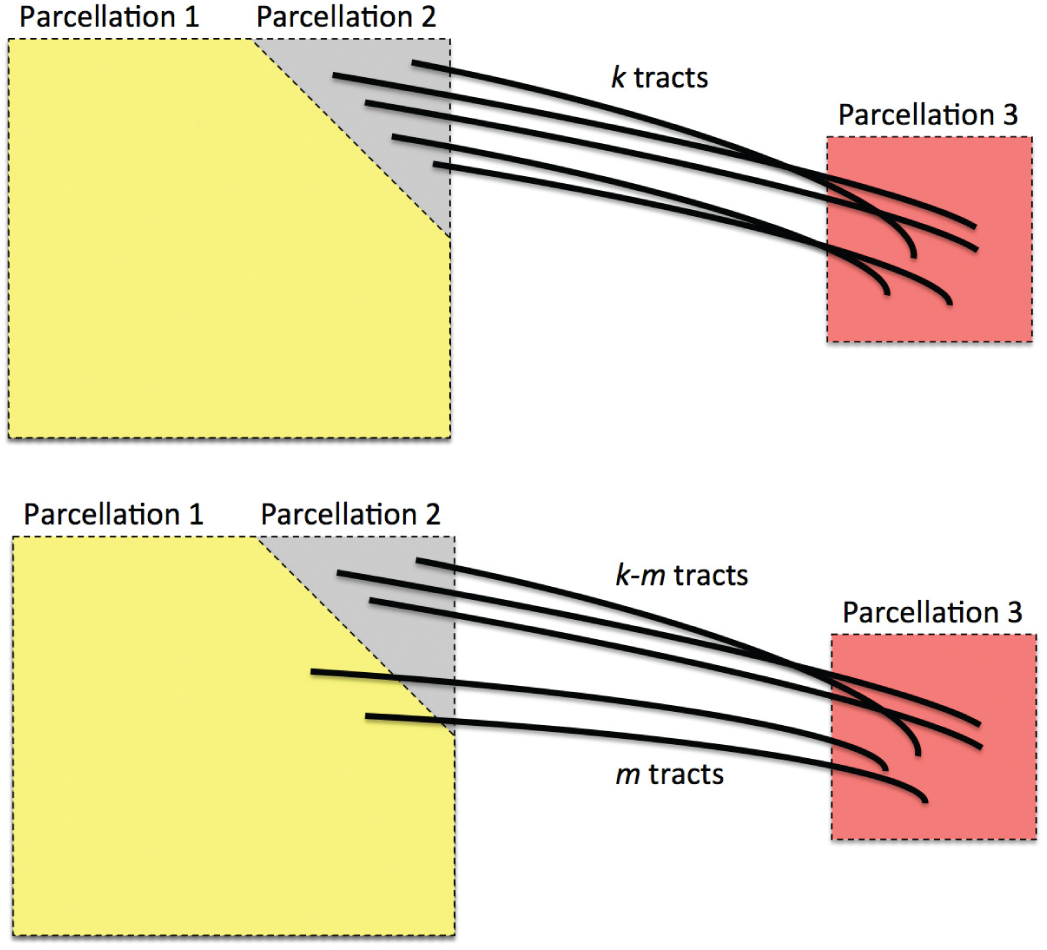
Schematic showing a possible tract mislabeling problem. *k* number of tracts are expected to pass through between parcellations 2 and 3 (top). However, *m* number of tracts might be mislabled to pass through parcellations 1 and 3 (bottom).

**Figure 8:**
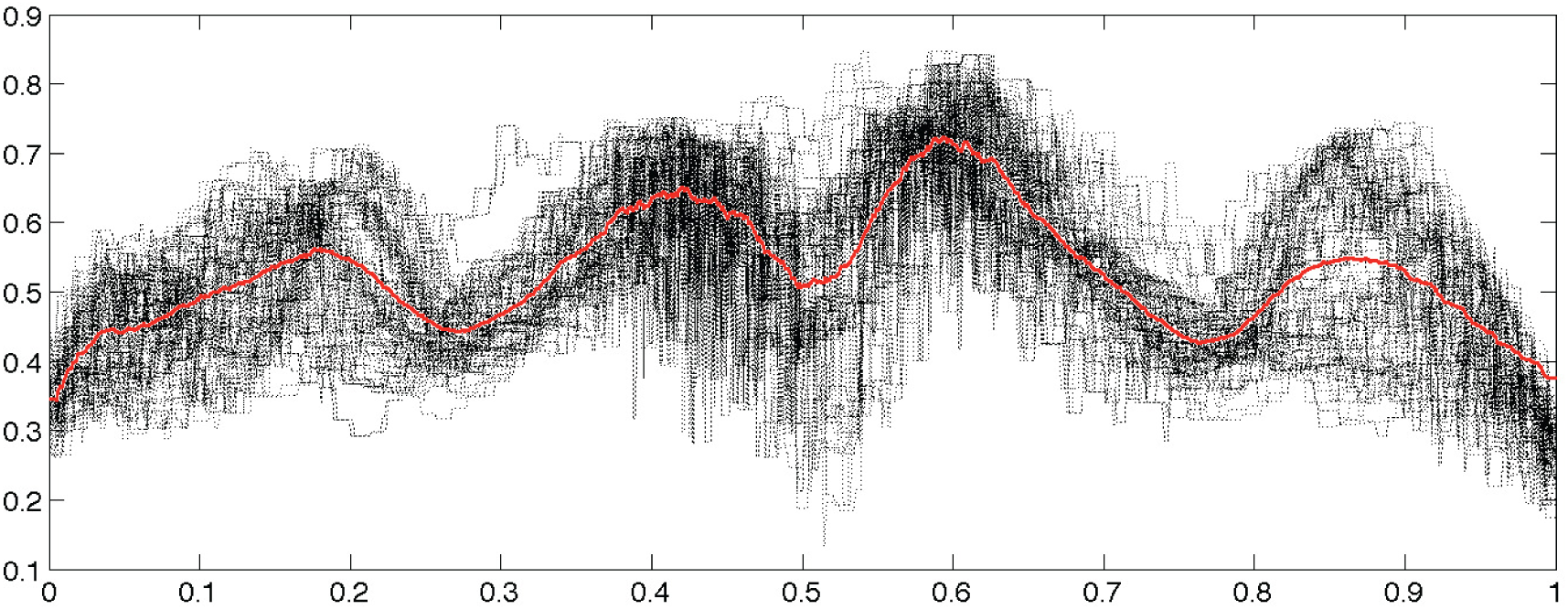
FA-values along 108 tracts between the left superior motor area (SMAG) to the right superior motor area (SMAD) displayed in Figure 1. The tracts are reparameterized between 0 and 1 from the left to right hemisphere. The red line is the average of FA-values of all the tracts

Suppose the *i*-th tract has length *L*_*i*_. The average tract length between the parcellations will be denoted as

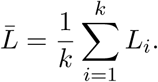

The resistance of the *i*-th tract is then given by

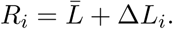

where 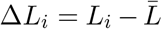 measures the difference from the mean. Subsequently, the expected total resistance is given by

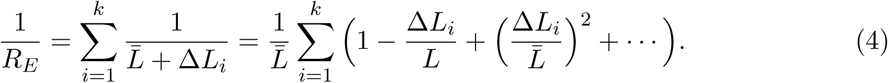

Since 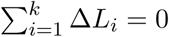, the expected total resistance is approximately

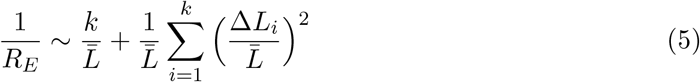

ignoring the cubic and other higher order terms. Similarly the observed resistance for *k* − *m* tracts is given by

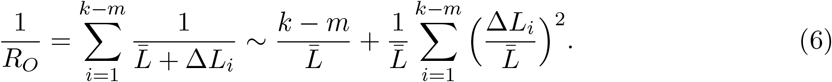

Hence the relative error of the new connectivity metric given by

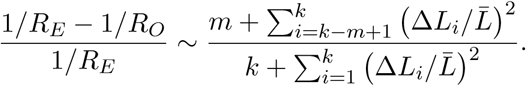

The terms 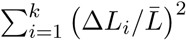 and 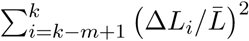 are sufficiently small relative and the relative error is approximately m/ *k* which is the relative error in the tract count based connectivity. Thus, we expect the error variability in the tract length based connectivity to scale proportionally to that of the tract count based method. So most likely all the methods will perform similarly in testing the connectivity differences at the edge level using the two-sample t-test since the t-test is scale invariant.

### 2.5 Degree Distributions

So far we have explored the issue of how to build weighted structural networks using three **DTI** features: tract length, counts and FA values. Here we show how to apply existing often used graph theoretical features on such networks. The graph theoretic features are too numerous to apply here. A review on other graph measures can be found in Bullmore and Sporns (2009). Probably the most often observed characteristic of complex network is scale-free property (Song et al., 2005). Thus, we will mainly explore if structural brain networks also follow scale-free among others using node degrees and degree distributions (Bullmore and Sporns, 2009).

The *degree distribution P* (*k*), probability distribution of the number of edges *k* in each node, can be represented by a power-law with a degree exponent *γ* usually in the range 2<*γ*<3 for diverse networks (Bullmore and Sporns, 2009; Song et al., 2005):

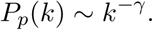

Such networks exhibits gradual decay of tail regions (heavy tail) and are said to be *scale-free*. In a scale-free network, a few hub nodes hold together many nodes while in a random network, there are no highly connected hub nodes. The smaller the value of *γ*, the more important the contribution of the hubs in the network.

Few previous studies have shown that the human brain network is not scale-free (Gong et al., 2009; Hagmann et al., 2008; Zalesky et al., 2010). Hagmann et al. (2008) reported that degree decayed exponentially, i.e.,

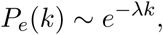

where *λ* is the rate of decay (Fornito et al., 2016). The smaller the value of *λ*, the more important the contribution of the hubs in the network.

Gong et al. (2009) and Zalesky et al. (2010) found the degree decayed in a heavy-tailed manner following an exponentially truncated power law

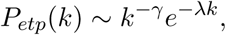

where 1*/λ* is the cut-off degree at which the power-law transitions to an exponential decay (Fornito et al., 2016). This is a more complicated model than the previous two models.

Directly estimating the parameters from the empirical distribution is challenging due to small sample size in the tail region. This is probably one of reasons we have conflicting results. To avoid the issue of sparse sampling in the tail region, the parameters are estimated from cumulative distribution functions (CDF) that accumulate the probability from low to high degrees and reduce the effect of noise in the tail region. For the exponentially truncated power law, for instance, the two parameters *γ, λ* are estimated by minimizing the sum of squared errors (SSE) using the L2-norm between theoretical CDF *F*_*etp*_ and empirical CDF 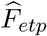:

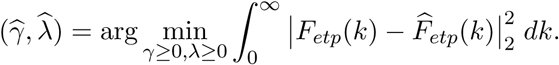

The estimated best model fit can be further used to compare the model fits among the three models. Many existing literature on graph theory features mainly deal with the issue of determining if the brain network follows one of the above laws (Fornito et al., 2016; Gong et al., 2009; Hagmann et al., 2008; Zalesky et al., 2010). However, such model fit was not often used for actual group level statistical analysis. Given two groups as in our study, we can fit one specific power model for each group. For instance, the exponential decay model can be fitted for each group separately and obtain different parameter estimates. Then one may test if the parameters are statistically different between the groups.

## 3 Results

### 3.1 Node degrees

Based on the three network construction methods, we computed the connectivity matrices. The connectivity matrices are binarized in such a way that any non-zero edges are assigned value one. This results in the adjacency matrices. Since the three connectivity methods only differ in the strength of the connectivity, the zero entries of the connectivity matrices exactly correspond across three connectivity matrices. Thus, the three adjacency matrices are identical. The degrees are then computed by summing the rows of the adjacency matrices. Figure 9 shows the mean node degrees for all 116 nodes for each group. The edges are the mean of the adjacency matrices above 0.5. We tested if there is any group differences in node degrees using the two-sample *t*-test (controls - maltreated). The maximum and minimum *t*-statistics are 2.95 (*p*-value = 0.0024) and -2.08 (*p*-value = 0.021). However, none of the nodes show any statistical significance after accounting for multiple comparisons using the false discovery rate (FDR) correction at 0.05.

**Figure 9:**
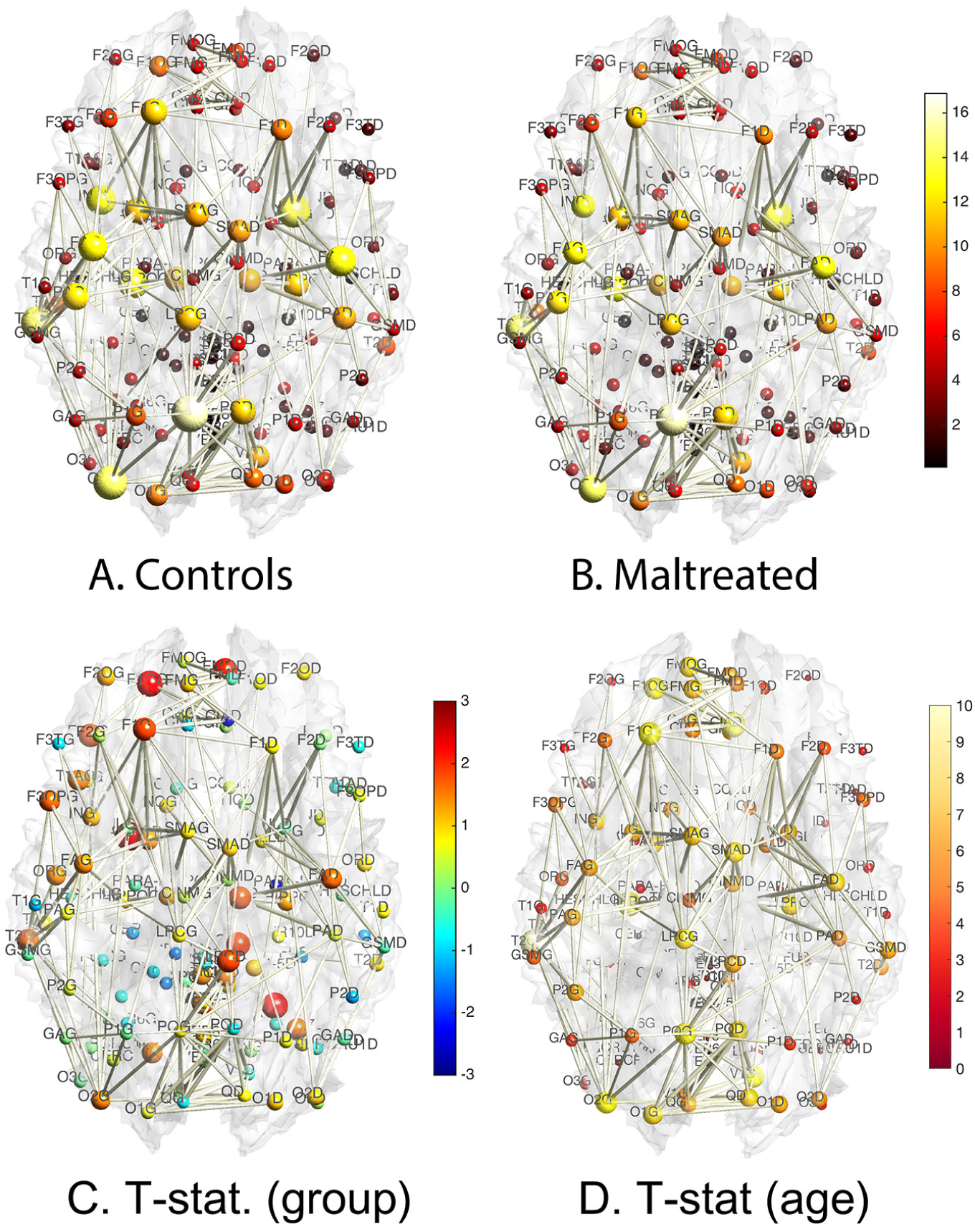
A. B. The node size and color correspond to the mean degree in each group. The edges are the average of the mean degrees of the two nodes threshoded at 0.5. There are consistent degree patterns across the groups although there are local differences. C. The two-sample *t*-statistic of the degree differences (controls - maltreated). There is no statistically significant node after the false discovery rate (FDR) correction at 0.05 level. D. *t*-statistic of age effect while accounting for sex and group variables. There is no significant reduction in node degree so only positive *t*-statistic values are shown. Most nodes passed FDR at 0.05 showing widespread connectivity increase in children.

We also looked at the effects of age and sex on node degrees. We tested the significance of sex while taking age and group as nuisance covariates in a general linear model. We did not detect any effect of sex at FDR at 0.05. We also tested the significance of group difference while accounting for age and sex. We did not detect any group difference at FDR at 0.05. However, we detected widespread age effect while accounting for sex and group variables at FDR at 0.05 (Figure 9-D). Almost all nodes (107 nodes out of total 116 nodes) showed statistically significant degree increases. There is no significant reduction in node degree so only positive *t*-statistic values are shown in the figure.

Due to the large variability associated with node degree, we were not able to differentiate subtle group differences locally although there is a consistent pattern of higher degree nodes in the controls. A more sophisticated approach is needed to see the degree differences. For this purpose, we investigated the degree distribution.

### 3.2 Degree distributions

We propose a two-step procedure for fitting node degree distribution. The underlying assumption of the two-step procedure is that each subject follows the same degree distribution law but with different parameters. In the first step, we need to determine which law the degree distribution follows at the group level. This is done by pooling every subject to increases the robustness of the fit. In the second step, we determine subject-specific parameter.

**Step 1.** This is a group level model fit. Figure 10-A shows the degree distributions of the combined subjects in each group. To determine if the degree distribution follows one of the three laws, we combined all the degrees across 54 subjects. Since high degree hub nodes are very rare, combining the node degrees across all subjects increases the robustness of the fit. This results in much more robust estimation of degree distribution. At the group level model fit, this is possible. The maltreated subjects had higher concentration in low degree nodes while the controls had high degree nodes. Thus the controls had much higher concentration of hub nodes. This pattern is also observed in the cumulative distribution functions (CDF) for each group (Figure 10-B). Again the maltreated subjects clearly show higher cumulative probability at lower degree.

**Figure 10:**
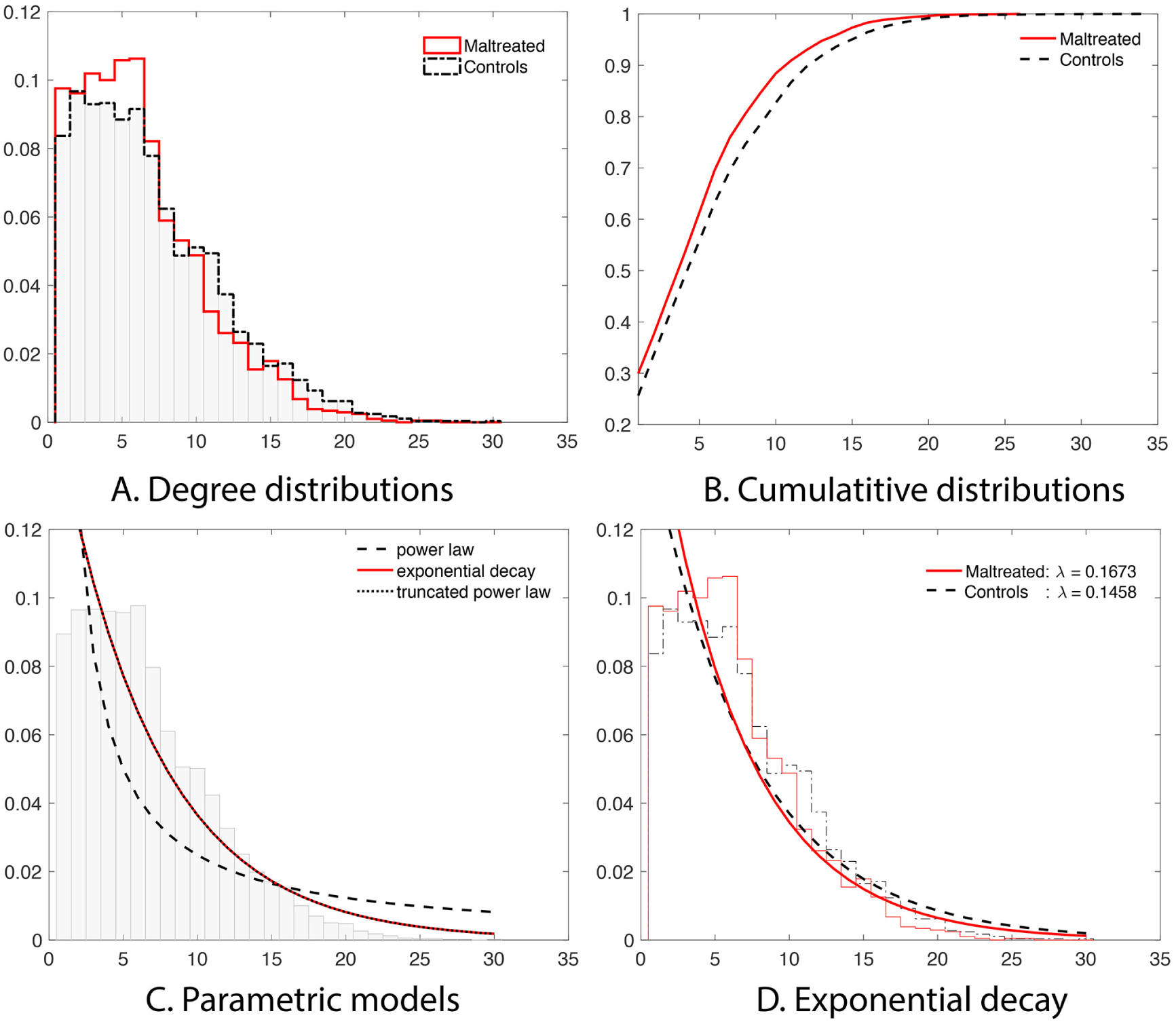
A. Degree distributions of all the subjects combined in each group. The normal controls have heavier tails for high degree nodes indicating more hub nodes. The maltreated children show much larger number of low degree nodes. B. The cumulative distribution functions (CDF) of all the subjects in each group. C. Three parametric model fit on the CDF of the combined 54 subjects. **D.** The exponential decay model is fitted in each group. The estimated parameters are significantly different (p-value<0.02).

The CDF of three laws were then fitted in the least squares fashion by minimizing the sum of squared errors (SSE) between the theoretical and empirical CDFs of the combined 54 subjects. The best fitting model for the power law was when *γ* = 0.2794, i.e.,

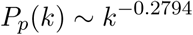

with SSE = 1.01. For the exponential decay law, the best model was when *λ* = 0.1500, i.e.,

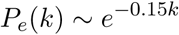

with SSE = 0.046. For the truncated power law, the best model was when *γ* = 0*, λ* = 0.1500, i.e.,

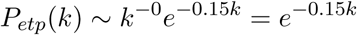

with SSE = 0.046. Note the best truncated power law collapses to the exponential decay when *γ* = 0. Thus, our data supports the exponential decay model as the degree distribution. Figure 10-C shows the three estimated models. Note the best truncated power law model is identical to the best exponential decay model for our data.

**Step 2.** This is the subject level model fit. We determined the degree distribution follows the exponential decay model at the group level. In the second step, we fitted the exponential decay model for each subject separately to see if there is any group difference in the estimated parameter. The estimated parameters were 0.1673 ± 0.0341 for the maltreated children and 0.1458±0.0314 for the controls. As expected, the controls showed a slightly heavier tail compared to that of the maltreated children (Figure 10-D). The two-sample *t*-test was performed on the parameter difference obtaining the significant result (*t*-stat. = 2.40, *p*-value<0.02). The controls had higher degree hub nodes, which are nodes with high degree of connections (Fornito et al., 2016).

Table 2 shows the list of 13 most connected nodes in the combined group in decreasing order. The numbers are the average node degrees in each group. We combined all the subjects in the two groups and computed the mean degree for each node. Then selected 13 highest mean degree nodes. All the 13 most connected nodes showed higher degree values in the controls without an exception.

**Table 2:** 13 most connected hubs in the combined group. They are sorted in the descending order of the degree. The controls have more connections without any an exception.

| Label | Parcellation Name | Combined | Controls | Maltreated |
| --- | --- | --- | --- | --- |
| PQG | Precuneus-L | 16.11 | 16.87 | 15.09 |
| NLD | Putamen-R | 14.96 | 15.26 | 14.57 |
| O2G | Occipital-Mid-L | 14.44 | 15.52 | 13.00 |
| T2G | Temporal-Mid-L | 14.30 | 15.16 | 13.13 |
| HIPPOG | Hippocampus-L | 13.15 | 13.94 | 12.09 |
| FAD | Precentral-R | 12.85 | 14.00 | 11.30 |
| ING | Insula-L | 12.56 | 13.61 | 11.13 |
| FAG | Precentral-L | 12.43 | 13.45 | 11.04 |
| PQD | Precuneus-R | 12.00 | 12.03 | 11.96 |
| PAG | Postcentral-L | 11.89 | 12.52 | 11.04 |
| NLG | Putamen-L | 11.39 | 11.68 | 11.00 |
| F1G | Frontal-Sup-L | 11.22 | 12.13 | 10.00 |
| HIPPOD | Hippocampus-R | 11.15 | 11.90 | 10.13 |

### 3.3 Inference on connectivity strength

Based on the tract counts, lengths and FA-values, the connectivity matrices were computed. Instead of thresholding the connectivity matrices, we looked at if there is any network differences at the edge level. Given 116 parcellations, there are total (116 *·* 115)*/*2 = 6670 possible edges connecting them. If all edges are connected, it forms a complete graph. However, few AAL parcellations are directly connected to each other. For our study, there are in average 1813 edges connecting across 116 regions. This reduces the number of tests in multiple comparisons as well in performing localized node-level statistical inference. Figure 5 shows the mean connectivity for first 32 nodes for the three methods. Figure 6 shows the graph representation of the connectivity matrices in each group.

We determined the statistical significance of the mean connectivity difference between the groups by performing the two-sample *t*-test (maltreated − controls). Only the connections that have *p*-value less than 0.05 (uncorrected) are shown in Figure 11 for all three methods (Figure 11-A to C). For the tract count method, max. *t*-stat. = 2.82 (*p*-value = 0.0034) and min. *t*-stat. = −2.92 (*p*-value = 0.0026). For the length-based method, max. *t*-stat. = 3.57 (*p*-value = 0.0004) and min. *t*-stat. = −3.18 (*p*-value = 0.0012). For the length-based method with FA, max. *t*-stat. = 3.45 (*p*-value = 0.0006) and min. *t*-stat. = −3.15 (*p*-value = 0.0014). Although there are major similarities between the three methods, no edge passed the multiple comparisons correction using FDR at 0.05. Even at FDR at 0.1 level, no edge was detected.

**Figure 11:**
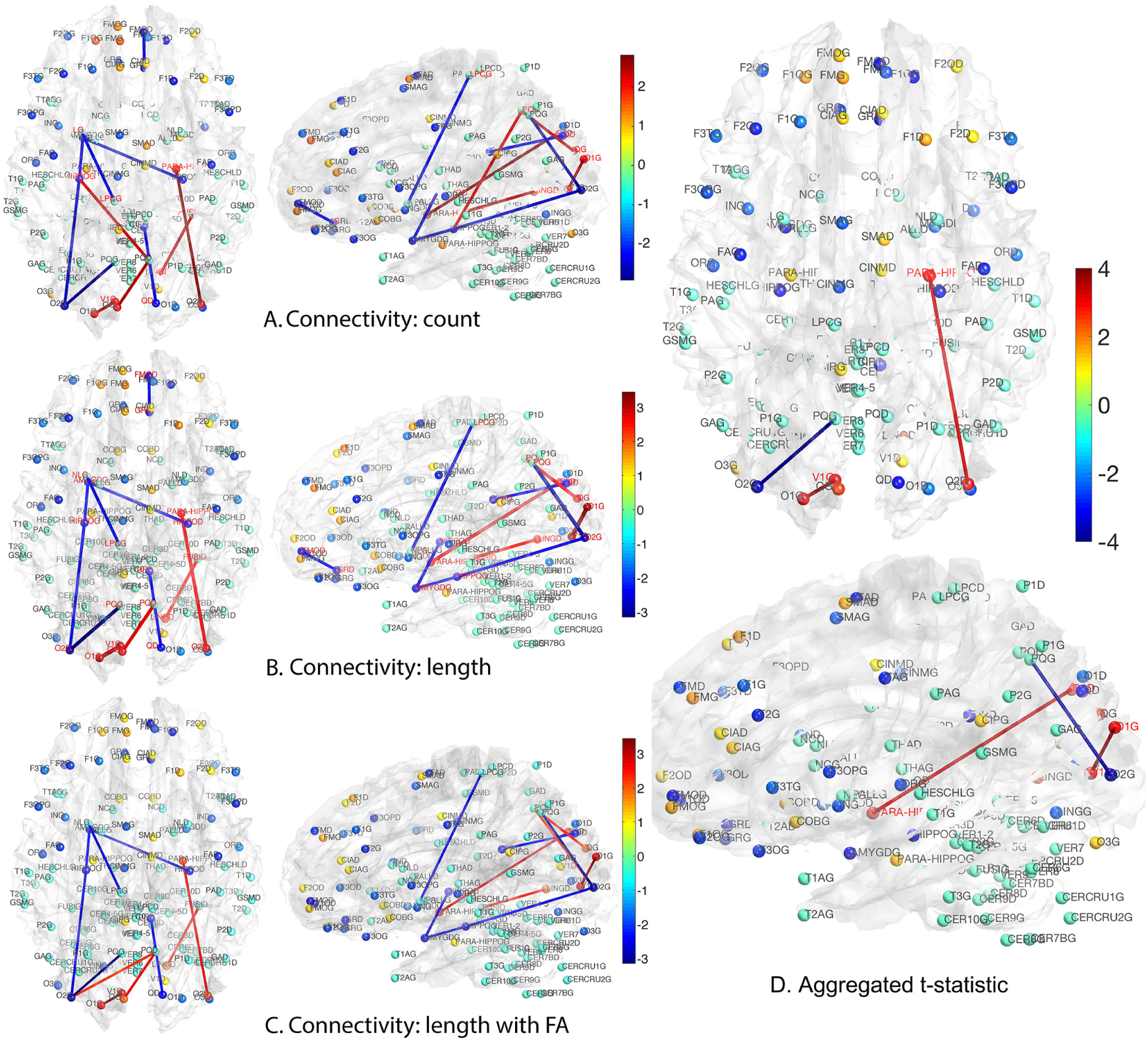
A.-C. *t*-statistic results (maltreated - controls) for three different connectivity methods. Only the connections at the *p*-value less than 0.01 (uncorrected) are shown. None of the edges pass FDR even at 0.1. All three methods give similar results showing the robustness and consistency. D. The three *t*-statistic maps are aggregated to form a single *t*-statistic. None of edges pass FDR at 0.05. Only edges passing FDR at 0.1 are shown.

Although there is no signal detected using the three methods after multiple comparisons correction, they are all giving the similar connectivity maps. Thus, we propose to integrate the three maps, by constructing the summary statistics map that aggregate the three *t*-statistics maps. We propose to use the sum of *t*-statistics that have been often used to aggregate multiple studies and samples and in meta analysis (Fan et al., 2004; Reimold et al., 2006). Previously, independently distributed *t*-statistics were used but the approach can be extended to dependent *t*-statistics that accounts for a more accurate variance estimate (Billingsley, 1995).

Suppose a collection of possibly dependent *t*-statistic maps *t*^1^,…,*t^n^* is given. We assume the degrees of the freedom (d.f.) of each *t*-statistic map is sufficiently large, i.e., d.f. ≥ 30. In our study, three *t*−statistic maps have 52 as the degrees of freedom. *t*-statistics for large degrees of freedom is very close to standard normal, i.e., *N* (0, 1). For *n* identically distributed possibly dependent *t*-statistics *t*^1^,…,*t^n^*, the variance of the 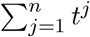 is approximately given by

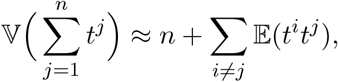

where 𝔼(*t^i^t^j^*) is the correlation between *t^i^* and *t^j^*. We used the fact 𝔼*t^j^* = 0. Then, we have the aggregated statistic *T* given by

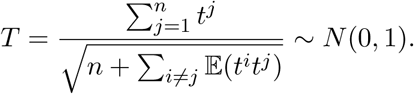

If the statistics *t^j^* are all independent, since *t^j^* are close to standard normal, 𝔼(*t^i^t^j^*) ≈ 0. The dependency increases the variance estimate and reduces the aggregated statistic value. Unfortunately, it may be difficult to estimate the correlation directly since only one *t*-statistic map is available for each *t^j^*. For our study, three dependent *t*-statistics maps are available. Thus, the variance is bounded between 3 = 1.73^2^ and 6 = 2.45^2^, and the aggregated *t*-statistic value is bounded between 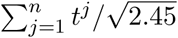 and 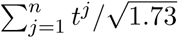. Unfortunately, it may not be easy to estimate the correlations directly since only one *t*-statistic map is available for each *t^j^*. In this study, 𝔼(*t^i^t*^*j*^) is empirically estimated by pooling over the entries of *t*-statistic maps *t*^*i*^ and *t^j^*. The result of the aggregated *t*−statistic map is given in Figure 11-D. Max. *t*−stat. = 3.96 (*p*-value = 3.76 × 10−5) and min. *t*-stat. = −3.78 (*p*-value = 7.73 × 10−5). Only the edges that passed FDR at 0.1 are shown but none of them passed FDR at 0.05.

We looked at the effects of age and sex in the analysis. We tested significance of sex by taking age and group as nuisance covariates in a general linear model. We did not detect any effect of sex in all three methods as well as the aggregated *t*-statistic at FDR at 0.05. We also tested the significance of group difference by taking age and sex as nuisance covariates. We did not detect any group difference in all three methods and the aggregated *t*-statistic at FDR at 0.05. However, we did detect significant and widespread age effect while accounting for sex and group variables in all three methods and the aggregated *t*-statistic even at very stringent FDR at 0.01 (Figure 12). Many major connections seem to show significant increase in connectivity strength. There is no significant reduction in connectivity so only positive *t*-statistic values are shown in the figure.

**Figure 12:**
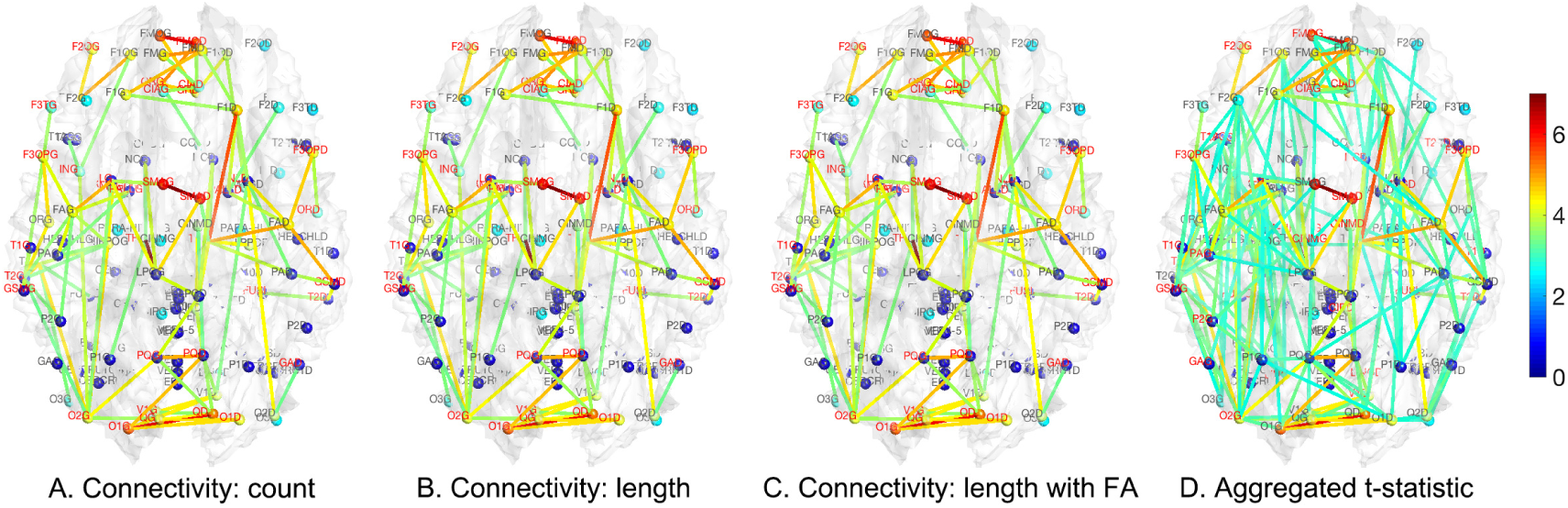
*t*-statistic map of age effect while accounting for gender and group variables. Only the significant edges thresholded at FDR at 0.01 are shown. There is no significant reduction in connectivity so only positive *t*-statistic values are shown. All three methods show consistent and widespread age effect. The aggregated *t*-statistic tend to enhance weak signal that are consistently presented in all three methods.

## 4 Conclusions & Discussion

Here, we presented a new integrative connectivity model for DTI, investigating three different DTI connectivity features: tract count, length and its FA values. The proposed integrative method can model the length of tracts as the resistance in the electrical circuit. The model can also incorporate FA-values into the model. The electrical circuit models without and with FA-values are then compared against the popular tract count method in characterizing connectivity in the maltreated children. The connectivity was quantified both at the node and edge levels.

Linking graphs to electric networks is not a new idea. In Doyle and Snell (1984), graph edge weights between nodes *i* and *j* are modeled as conductance *d*_*ij*_, which is the inverse of resistance, i.e.,

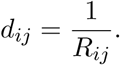

Then the a random walk on the network is defined as a Markov chain with transition probability

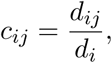

where 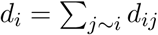, the sum of all conductance of edges connecting nod *i*. The denominator has the effect of normalizing the conductance into a proper probability measure. The transition probability matrix *C* = (*c*_*ij*_) then can be used as a connectivity matrix. However, this is not the only way to normalize the conductance *d*_*ij*_ to make it as a probability. Instead of normalizing locally at each node (a column or row wise normalization), one alternative is to normalize globally by dividing the numerator by either max_*ij*_*d*_*ij*_ or 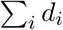. Since the statistical analyses we used are scale invariant, we should obtain exactly the same results if such global normalization was performed.

The proposed model is more of an analogy based on an existing electrical circuit model. DTI-based brain networks at the macroscopic level may not follow the physical laws of electrons passing through a wire. It was not our intention to mimic the actual electrons in neuronal fibers using electrical circuits. We tried to come up with a mathematical model that can incorporate tract counts, lengths and FA-values into a single connectivity metric and the electrical circuit model seems to be a reasonable model to use. There can be many possible combination of methods to construct connectivity in DTI. Unless we test for every possible combinations, it may be difficult to address the problem. More research is needed to identify the most representative connectivity measure that incorporates various structural properties obtained from DTI. In this paper, we showed that within some degree, normalizations may not really matter to the final statistics maps and all three methods gave more or less the same results.

To demonstrate the proposed method gives reasonably consistent results against the existing tract count method, we performed a theoretical error analysis in Section 2.4 and have shown that the error variability in the resistance method scales proportionally to that of the tract count method. This guarantees the resulting statistical maps are more or less similar to each other. Since the statistics are scale invariant, it is expected we will have similar connectivity maps as shown in Figure 11 and 12, where A (count), B (length) and C (length with FA) are almost identical. Denote *t*^1^, *t*^2^ and *t*^3^ as the *t*-statistics maps from the tract count, length-based and FA-value based methods respectively. For the two sample *t*-test on group variable, we have *corr*(*t*^1^, *t*^2^) = 0.99, *corr*(*t*^2^, *t*^3^) = 1.00, *corr*(*t*^1^, *t*^3^) = 0.99. For *t*-statistic on age while accounting for group and sex is *corr*(*t*^1^, *t*^2^) = 0.99, *corr*(*t*^2^, *t*^3^) = 1.00, *corr*(*t*^1^, *t*^3^) = 1.00. So even though three different connectivity metrics are used, we end up with almost identical connectivity maps at the end.

We determined the structural brain network follows the exponential decay law. However, Gong et al. (2009) reported the brain to follow the more complicated truncated exponential power law by comparing the goodness-of-fit of the model using *R*^2^-value. The truncated exponential power law has two parameters while the exponential decay law has one parameter. The exponential decay law is a special case of the truncated exponential power law when *γ* = 0. Thus, the truncated exponential power law will fit better than the exponential decay law for any data. However, in our data, the *γ* = 0 and the truncated exponential power law collapsed to the exponential decay law. For other datasets, in general, the truncated exponential power law will fit better.

The group level statistical results show very similar network differences although they did not identify any significant edges using FDR at 0.05. Although there are few pervious connectivity studies that incorporated tract lengths and FA-values (Skudlarski et al., 2008; Kim et al., 2015), we found the inclusion of additional DTI features into connectivity model do not really change the final statistical results much. All these DTI features are likely linearly scaling up. Since most statistics are scale invariant, we may obtain similar results. To integrate the similar but different statistical maps, we employed a meta-analytic framework to aggregate the results into a single statistical map. This method seemed to boost signal when there are weak signals, i.e., low *t*-statistic values, but that are consistently presented.

## Acknowledgements

This work was supported by NIH Research Grants MH61285, MH68858, MH84051, UL1TR000427 and Brain Inititative Grant EB02285.

